# Discarded diversity: Novel megaphages, auxiliary metabolic genes, and virally encoded CRISPR-Cas systems in landfills

**DOI:** 10.1101/2024.05.30.596742

**Authors:** Nikhil A. George, Zhichao Zhou, Karthik Anantharaman, Laura A. Hug

## Abstract

**Background:** Viruses are the most abundant microbial entity on the planet, impacting microbial community structure and ecosystem services. Despite outnumbering Bacteria and Archaea by an order of magnitude, viruses have been comparatively underrepresented in reference databases. Metagenomic examinations have illustrated that viruses of Bacteria and Archaea have been specifically understudied in engineered environments. Here we employed metagenomic and computational biology methods to examine the diversity, host interactions, and genetic systems of viruses predicted from 27 samples taken from three municipal landfills across North America.

**Results:** We identified numerous viruses that are not represented in reference databases, including the third largest bacteriophage genome identified to date (∼678 kbp), and note a cosmopolitan diversity of viruses in landfills that are distinct from viromes in other systems. Host-virus interactions were examined via host CRISPR spacer to viral protospacer mapping which captured hyper-targeted viral populations and six viral populations predicted to infect across multiple phyla. Virally-encoded auxiliary metabolic genes (AMGs) were identified with the potential to augment hosts’ methane, sulfur, and contaminant degradation metabolisms, including AMGs not previously reported in literature. CRISPR arrays and CRISPR-Cas systems were identified from predicted viral genomes, including the two largest bacteriophage genomes to contain these genetic features. Some virally encoded Cas effector proteins appear distinct relative to previously reported Cas systems and are interesting targets for potential genome editing tools.

**Conclusions:** Our observations indicate landfills, as heterogeneous contaminated sites with unique selective pressures, are key locations for diverse viruses and atypical virus-host dynamics.

## Background

In one of the largest examinations of earth’s virosphere to date, it was noted that samples from engineered sites have poorly characterized viral diversity [1]. Two recent studies of wastewater treatment plants confirmed this, with >98% of detected viral contigs unmatched to a representative subset of the IMG/VR database in one [2] and >99% unmatched to the NCBI virus database in the other [3]. Landfills, similar to wastewater treatment plants, are highly engineered, polluted and environmentally significant systems [4–6]. Examinations of CRISPR array variation and spacer conservation have been performed across a variety of sample types [7], including in wastewater [3], but these examinations have not been performed with samples from municipal waste sites.

Bacteriophages are predicted to be quite host specific, but recent studies have suggested the presence of phages with diverse host ranges in different environments. In research on both isolated phages [8] and those identified *in silico*, bacteriophages have been shown or predicted to infect across distinct bacterial phyla [1,9–11] and domains of life [2,8,11]. While the mechanisms of viral infection across organisms from different phyla or domains are likely very complex and difficult to resolve without wet-lab experiments, the potential presence of such viruses with large host ranges and their isolation can be important for biotechnological applications including phage therapy. There are multiple computational methods to predict microbial hosts of viruses [12,13], with CRISPR spacer to viral protospacer matching having relatively low sensitivity, but high specificity [12]. In municipal waste systems, viruses that infect bacteria and archaea are predicted to have diverse infection dynamics with their hosts based on CRISPR spacer to virus protospacer matching [9]. The open questions remain of whether the notable virus-host interactions observed in a single municipal waste site, such as hyper-targeted viral elements, interviral conflicts, and putative cross-phylum-infecting phages [9] are consistent across other municipal waste sites, and whether these interactions are underlain by conserved CRISPR array architectures across geographically distant sites.

Viruses of bacteria and archaea have genomes that are variable in size, but are canonically considered short, efficient genomic elements constrained by their requirement to package into capsids for infection. Recently, genomic approaches have identified an increasing number of phage genomes ≥200 kbp in size, where phages with genomes ≥200 kbp are referred to as jumbophages and phages with genomes ≥500 kbp are referred to as megaphages [1,14,15]. The current record length for a circularized megaphage genome is 735 kbp [16]. These less-streamlined phage genomes have more coding space, and have been shown to carry genes encoding tRNAs and associated enzymes, other translation machinery, auxiliary metabolic genes (AMGs), and CRISPR-Cas systems [16].

During infections, viruses may augment or modify the metabolism of their hosts through AMGs [17–21], including contaminant modification and degradation. Recent research has identified viruses encoding arsenic resistance and transporter genes in arsenic-contaminated paddy soils [22], and showed viruses aiding their hosts’ survival in organochlorine contaminated soils with AMGs involved in organochlorine degradation [23,24]. This phenomenon of contaminant modification/degradation genes being provisioned by viruses to their hosts has yet to be examined in municipal waste sites, which are highly and heterogeneously contaminated systems.

Despite the widespread adoption of CRISPR-Cas-based methods for genome editing and other biotechnological tools in both academic and industrial labs, the technology still has limitations [25] . Discovery of new, divergent Cas proteins and CRISPR-Cas system assemblages is of interest to address these limitations, in hopes of broadening the current application space [25] . Virus-encoded CRISPR arrays and CRISPR-Cas systems have been identified from a variety of environments [9,16, 26–33] and are involved in crippling host viral defense systems [28]; regulating host transcription and translation[16]; CRISPR-Cas system inhibition [31]; and interviral conflicts [9,31–32] . Recent discoveries of some of the most streamlined CRISPR-Cas systems from viruses [16,33,34] established viruses as a potential hot spot for new CRISPR-Cas systems with biotechnological value.

Here we examine the viral diversity within three municipal landfills distributed across North America, examining host-virus dynamics, CRISPR array conservation across geographic distance, and the unique diversity of the landfill virome. We identify new lineages within the jumbo and megaphage viral pantheon and characterize AMGs that the viral fraction is contributing to the landfill microbial community. Finally, we expand the examination of virally encoded CRISPR-Cas systems to municipal landfill microbial communities, identifying several streamlined CRISPR-Cas systems as potential targets for developing efficient biotechnological tools.

## Results

### Sampling sites and metagenomes

This study makes use of 27 metagenomes that were sequenced from samples from three distinct landfills. Fourteen metagenomes were sequenced from an active municipal landfill and adjacent aquifer in Southern Ontario (SO), sampled twice one year apart (July 2016 [SO 2016] and October 2017 [SO 2017]) and described previously [9,35]. Nine metagenomes were from a northeastern United States active municipal landfill (NEUS) in which nine distinct cells, units in which waste is held within the landfill, were sampled in February 2019 as described previously [36]. At the time of sampling, the oldest cell had received waste from 1980 to 1982 (39 years of waste decomposition time), whereas the youngest cell began receiving waste in 2014 (5 years of waste decomposition time). The final four metagenomes came from a closed landfill from Southern California that was sampled in June of 2019 (CA_2019). The Californian landfill was operational from the mid 1960s to the mid 1990s and received both municipal and hazardous waste. The landfill has been closed for over 20 years but remains under management for contaminant containment. From the three landfill sites, samples were taken from leachate wells, leachate collection cisterns, leachate-impacted groundwater wells, and, in one case, a leachate treatment facility (CA landfill, influent and biosolids fraction sampled) (Table S1).

### Viral proportion and genome sizes

Viral scaffold prediction was conducted by VirSorter2 (VS2) [37]. Viral genome quality was checked with CheckV [38], followed by clustering with CD-HIT for all sequences above 5 kb in length [39], which resulted in a total of 22,658 unique viral scaffolds in the SO 2016 samples (27 scaffolds ≥ 200 kbp). The SO 2017 samples had a total of 37,182 unique viral scaffolds (69 scaffolds ≥ 200 kbp), the NEUS samples had a total of 31,944 unique viral scaffolds (31 scaffolds ≥ 200 kbp), and the CA_2019 samples had a total of 5,419 unique viral scaffolds (11 scaffolds ≥ 200 kbp). Collectively, 97,203 viral scaffolds ≥ 5 kbp were identified (Table 1). Notably, seven of these viral scaffolds were predicted to be nearly complete or complete megaphage genomes as they were ≥ 500 kbp in length. Six out of seven of these sequences were predicted to be 100% complete according to CheckV’s identification of high-confidence Direct Terminal Repeats (DTRs) in the sequences, and all seven sequences were predicted to circularize by vRhyme (Table 2, [22]). Large terminase subunit genes, genes encoding part of the viral DNA packaging machinery, were identified in six out of seven putative megaphages, with five of these sequences placing within the Mahaphage clade of giant phages [16] in a phylogenetic tree (Figure 1), and one of them placing within an unclassified lineage under the most recent classification scheme for giant phages [16]. The two largest putative megaphage sequences were ∼641 kbp (NEUS_megaphage_1) and ∼678 kbp (SO_2017_megaphage_1) in size. The 678 kbp phage is, to our knowledge, the third largest phage genome reported to date, with the largest previously reported phage genomes being 634, 636, 642, and 735 kbp [16], 656 kbp [41], 660kbp [42], and 717 kbp [43].

**Figure 1:**
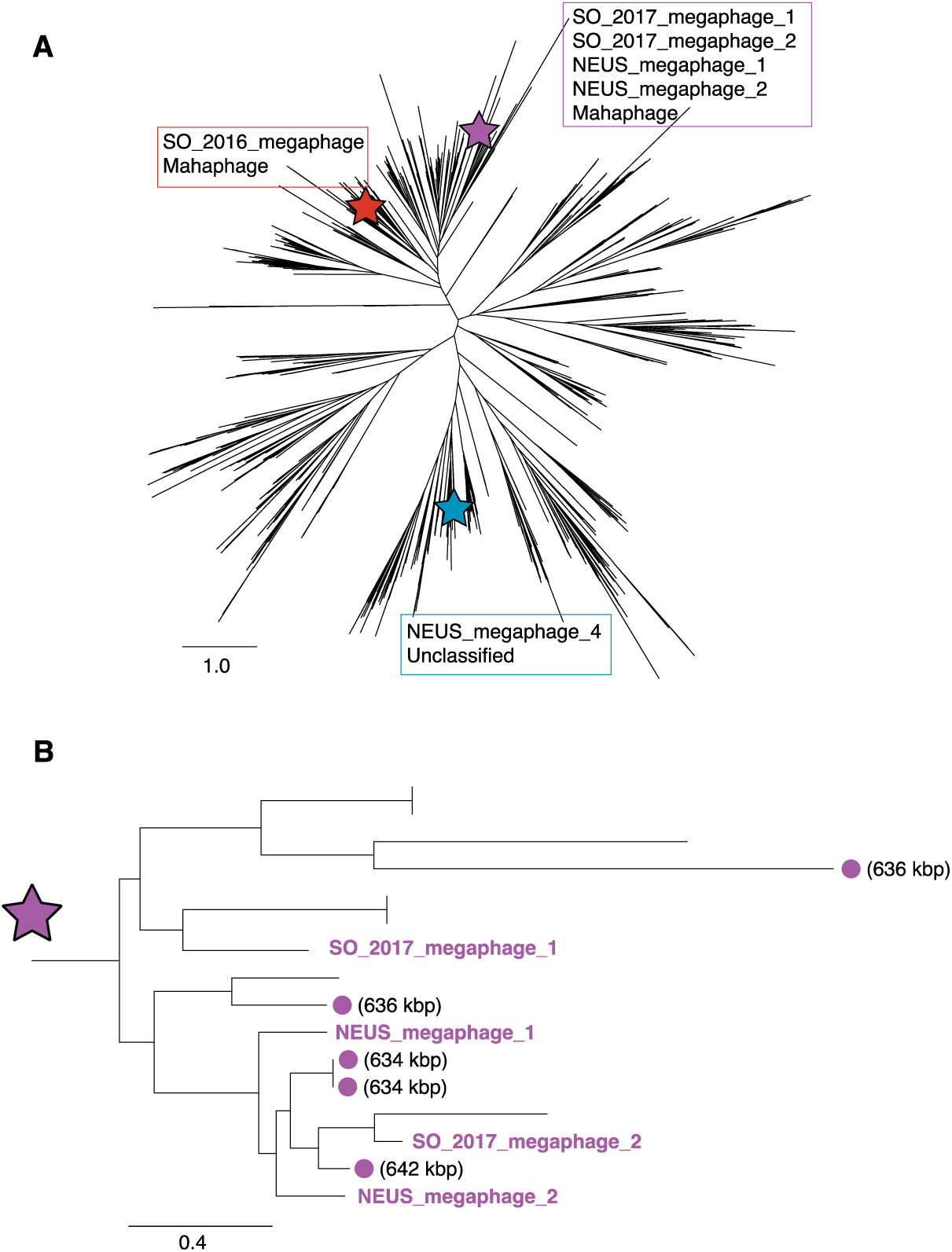
A: Phylogenetic placement of newly discovered megaphages. Phylogenetic reconstruction was based on an alignment of large terminase subunit encoding genes from this study (n=6) and those from a reference study (n=1,256, [16]). Stars indicate the clades in which megaphages identified in this study were placed [16]. B: Placement of four newly discovered megaphages in a clade specific to huge phages. References phages in this clade include some of the largest phages identified from a previous study [16]. The sizes of reference phage genomes are indicated if they are ≥600 kbp.

**Table 1:** Summary statistics for metagenomes, MAGs, and viral scaffolds from the sampling sites.

| Site | # of<br>Metagenomes | Read count | Scaffolds<br>≥5 kbp | MAGs<sup>1</sup> | viral<br>scaffolds<br>≥5 kbp<sup>2</sup> | %<br>viral<sup>3</sup> |
| --- | --- | --- | --- | --- | --- | --- |
| SO_2016 | 6 | 1,135,023,824 | 190,754 | 688 | 23,893 | 12.43 |
| SO_2017 | 8 | 1,822,382,124 | 269,004 | 1,064 | 39,612 | 13.53 |
| NEUS_2019 | 9 | 1,869,347,618 | 530,926 | 1,881 | 36,453 | 6.66 |
| CA_2019 | 4 | 862,249,102 | 231,655 | 762 | 6,410 | 2.15 |

<sup>1</sup> – Quality filtered bins with >70% completion, <10% contamination
<sup>2</sup> – viral scaffolds detected (prior to CD-HIT clustering)
<sup>3</sup> – % of total nucleotides in assembled scaffolds ≥5 kbp predicted to be of viral origin

**Table 2:**
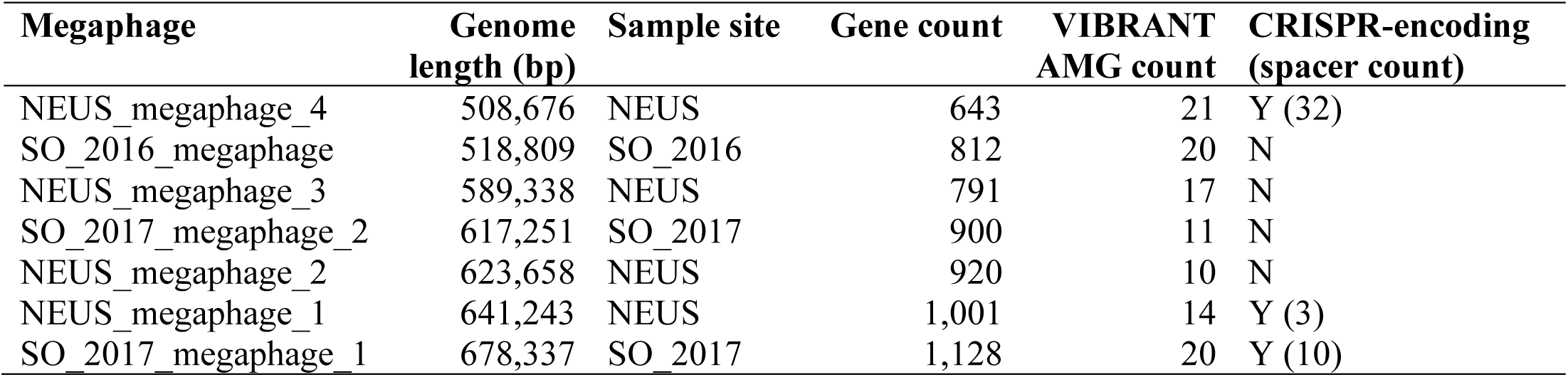
Genome statistics for complete megaphages identified in this study.

| **Megaphage** | **Genome**<br>**length (bp)** | **Sample site** | **Gene count** | **VIBRANT**<br>**AMG count** | **CRISPR-encoding**<br>**(spacer count)** |
| --- | --- | --- | --- | --- | --- |
| NEUS_megaphage_4 | 508,676 | NEUS | 643 | 21 | Y (32) |
| SO_2016_megaphage | 518,809 | SO_2016 | 812 | 20 | N |
| NEUS_megaphage_3 | 589,338 | NEUS | 791 | 17 | N |
| SO_2017_megaphage_2 | 617,251 | SO_2017 | 900 | 11 | N |
| NEUS_megaphage_2 | 623,658 | NEUS | 920 | 10 | N |
| NEUS_megaphage_1 | 641,243 | NEUS | 1,001 | 14 | Y (3) |
| SO_2017_megaphage_1 | 678,337 | SO_2017 | 1,128 | 20 | Y (10) |

### Genetic relatedness of viruses across ecosystems and viral taxonomy

We were interested in whether landfill viruses were conserved across sites, or across ecosystems. We used vConTACT2 [44] to determine the extent of relatedness between viral genomes in our study. vConTACT2 uses shared protein clusters (PCs) to delineate viral clusters (VCs) and was run against all ≥10 kbp viral elements (34,499) detected in our samples, along with a total of 45,523 viral elements from IMG/VR [45] detected in either wastewater samples (24,982) or groundwater samples (20,541), and the “Prokaryotic Viral Refseq version 85 with ICTV-only taxonomy” reference database (2,617 reference sequences) (Figure S1). While some similarity was observed between landfill viral elements and those from wastewater and groundwater, landfill viral elements frequently formed their own clusters with limited overlap with viral elements from other sites. We next used vConTACT2 to examine how similar viral communities were across the landfill environments (Figure S2). Compared to the gene-sharing network that included wastewater and groundwater viruses (Figure S1), the gene-sharing network including only landfill-associated viruses showed a much higher level of viral clustering (Figure S2). The broader environmental network contained 13,163 VCs and the landfill-only network contained only 6,463 VCs in a more strongly interconnected network. Using geNomad, 33,609 (∼97%) of our >= 10 kbp viral elements were classified to the class level as Caudoviricetes and only 573 (∼1.7%) were classified at either the order level (or family level, in cases where there was no assigned order).

### Host-virus connections within and across landfills

To identify hosts of viruses, we examined matches between CRISPR spacers to viral protospacers targeting both within a particular landfill site and across distinct landfills/times. SO_2016 showed the strongest targeting to NEUS_2019, having more spacer to protospacer matches than NEUS_2019 did to SO_2016, 327 vs. 262, despite NEUS_2019 having almost four times the number of spacers as SO_2016 (Table 3). SO_2017 showed strongest targeting to its preceding temporal dataset, SO_2016. NEUS_2019 spacers targeted viruses in its own dataset most strongly. Lastly, CA_2019 was targeted least often by spacers from other datasets. Despite having, at minimum, nearly four times fewer viral elements than any other dataset, CA_2019 spacer to protospacer matches was the second highest out of all 16 comparisons (853 connections, Table 3). All sample sites showed instances of hyper-targeted viral elements and/or vMAGs, which we previously defined as viral elements (or vMAGs) targeted at least 20 times by a single host’s complement of CRISPR spacers [9]. The maximum number of spacer-to-protospacer matches between a host MAG and a viral element (or vMAG) per dataset are 55 (SO_2016), 24 (SO_2017), 32 (CA_2019), and 22 (NEUS_2019). After curation, across all networks, a total of five vMAGs and one viral element were predicted to infect across distinct phyla (Table S2), with one vMAG from NEUS_2019 predicted to infect hosts from three distinct bacterial phyla: Bacteroidota, Cloacimonadota, and Firmicutes_B. The SO_2016 network was the only network that didn’t include putative cross-phylum infections. Lastly, considering the relatively low viral scaffold count from the CA_2019 dataset (Table 3), it showed relatively high host and viral interconnectivity in its network (Table 4).

**Table 3:** Host CRISPR spacer to viral protospacer targeting within and across municipal waste sites. Host CRISPR spacers and viral elements were not dereplicated for this analysis. VE = viral elements.

| Host targeting<br>(CRISPR spacer count) | # of spacer-protospacer matches (# VE) |  |  |  |
| --- | --- | --- | --- | --- |
|  | to SO_2016<br>(23,893 VE) | to SO_2017<br>(39,612 VE) | to NEUS_2019<br>(36,453 VE) | to CA_2019<br>(6,410 VE) |
| SO_2016 (5,446) | 266 | 186 | 327 | 12 |
| SO_2017 (12,578) | 461 | 451 | 443 | 21 |
| NEUS_2019 (20,236) | 262 | 654 | 1,595 | 79 |
| CA_2019 (9,299) | 119 | 39 | 121 | 853 |

**Table 4:** Virus-host networks summary. Virus-host interactions are based on host MAG CRISPR spacers and vMAG or viral element (depending on if element was binned) protospacer connections. Unique links = 1 MAG to 1 viral element, regardless of number of spacer-protospacer connections.

| Network | Host MAGs | Viral scaffolds or vMAGs | Unique links |
| --- | --- | --- | --- |
| SO_2016 | 13 | 20 | 22 |
| SO_2017 | 73 | 100 | 115 |
| NEUS_2019 | 169 | 206 | 321 |
| CA_2019 | 68 | 86 | 127 |

### CRISPR spacer conservation across waste sites

To assess the conservation of viral defenses across landfills, we next examined conservation of specific CRISPR spacers within and between landfills (Table 5). Generally, SO_2016 and SO_2017 had the most spacers in common, which was expected given they are temporal samples from the same location, and with overlapping sampling sites. The CA_2019 spacer set had very limited overlap with the other landfill sites.

**Table 5:** Shared CRISPR spacers between landfill datasets. Each dataset’s host spacers were dereplicated by CD-Hit at 100% identity for this analysis. Shared spacers required exact matches. Reverse complement matches were included in this analysis.

| **Dataset** | **Number of host spacers** | **Spacers shared with SO_2016** | **Spacers shared with SO_2017** | **Spacers shared with NEUS_2019** | **Spacers shared with CA_2019** |
| --- | --- | --- | --- | --- | --- |
| SO_2016 | 5,361 | N/A | 1,252 | 162 | 6 |
| SO_2017 | 11,812 | 1,252 | N/A | 174 | 10 |
| NEUS_2019 | 17,581 | 162 | 174 | N/A | 17 |
| CA_2019 | 7,223 | 6 | 10 | 17 | N/A |

### Viral auxiliary metabolic genes

We next assessed the contribution from viruses to community metabolic processes by predicting AMGs encoded on viral elements. Across all landfills, a total of 582 unique AMGs were predicted (Table S3), representing functions across all major cellular processes (Figure 2). AMG profiles across different landfills and compared to the IMG/VR dataset were quite similar, except for the CA_2019 AMGs (Figure 2). CA_2019 had relatively high counts of putative AMGs associated with lipid, amino acid, and aromatic compound metabolism, and comparatively low counts of AMGs associated with carbohydrate and terpenoids/polyketide metabolism. AMGs for processes that had not previously been associated with viruses or those that had very limited representation in literature were further manually curated to ensure they were not the first or final gene on a viral element, that they were flanked by viral genes or solely hypothetical proteins, and that no cellular organismal marker genes were present on the relevant elements. AMGs of interest that passed manual curation included genes predicted to be involved in carbohydrate metabolism, methane metabolism, and sulfur metabolism (Table 6). The CA_2019 dataset included an AMG predicted to have a role in degradation of organochlorine compounds, which has been observed in previous studies [23,24]. Numerous putative AMGs were also predicted in the putative megaphages - full details of VIBRANT-predicted AMGs encoded by these putative megaphages are available in Table S4.

**Figure 2:**
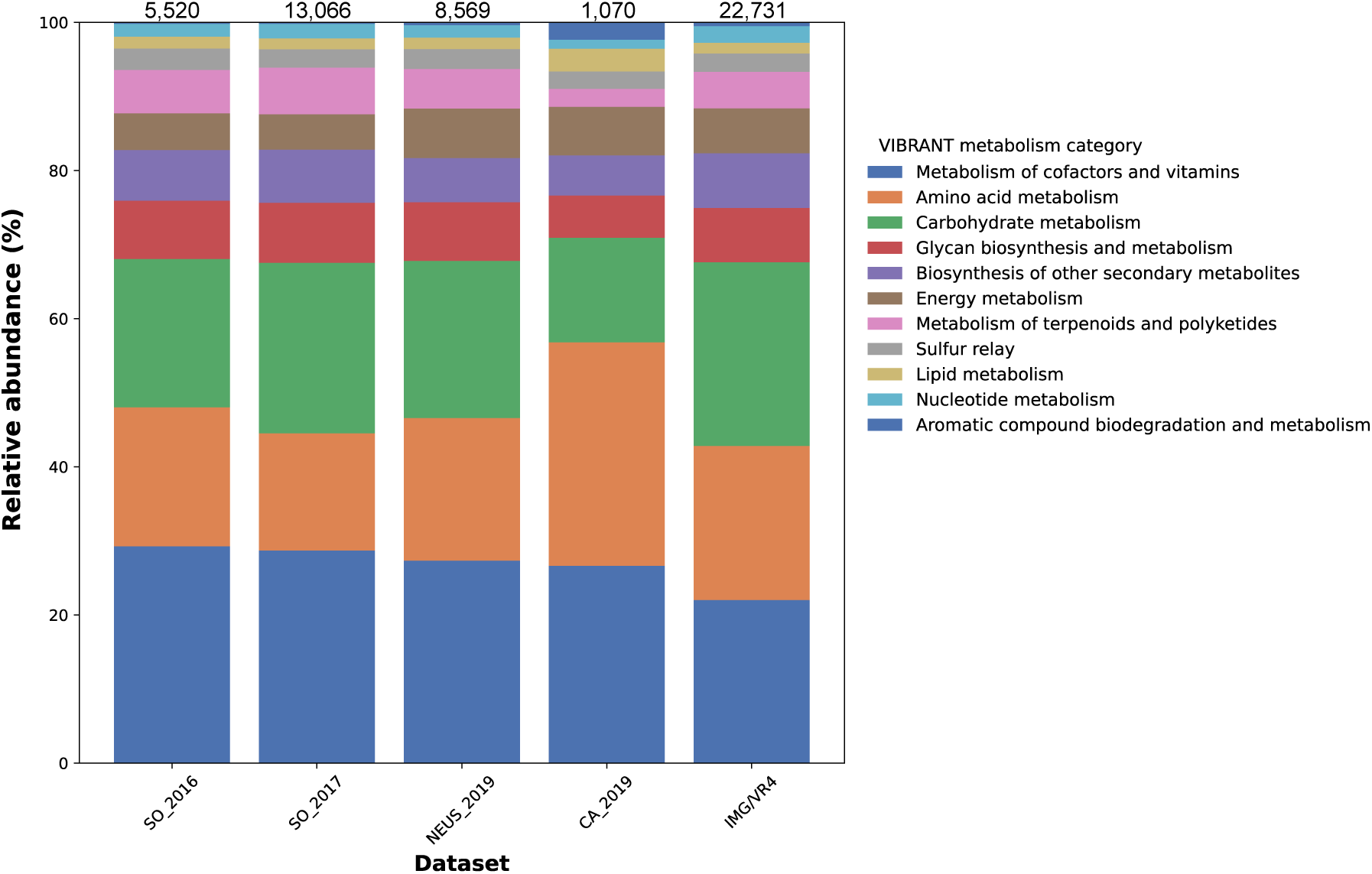
AMG metabolism category proportions across datasets. AMGs were detected using VIBRANT [46]. The IMG/VR dataset was used as a reference and was generated by randomly sampling 100,000 viruses from IMG/VR4 that were ≥5 kbp in size. The legend denotes AMG metabolism categories assigned by VIBRANT. Counts of viral elements ≥5 kbp included in the AMG search are as follows: SO_2016 - 23,893, SO_2017 - 39,612, NEUS_2019 - 36,453, CA_2019 - 6,410. Counts of predicted AMGs per dataset are reported at the top of each bar. The 22,731 AMGs were predicted from the IMG/VR dataset, some of which may not have been previously reported.

**Table 6:** Notable putative auxiliary metabolic genes encoded by viruses in the landfill datasets. Entries marked with an * represent functions also observed in 100k randomly selected IMG/VR prokaryotic viruses, but which are not widely reported in the literature. Functions are presented in alphabetical order according to AMG name. Highest confidence AMGs are marked with an ^ and explored further with phylogenetic trees.

| AMG name | Sites: #<br>instances<br>per site | AMG function |
| --- | --- | --- |
| *aprB*; adenylylsulfate reductase, subunit B [EC:1.8.99.2] | SO_2017: 1 | Sulfur metabolism |
| *dehH*; haloacetate dehalogenase [EC:3.8.1.3] | CA:1 | Chlorocyclohexane and chlorobenzene degradation, Chloroalkane and chloroalkene degradation |
| ^E3.2.1.1A; alpha-amylase [EC:3.2.1.1] | NEUS: 1 | Polysaccharide metabolism |
| ^*frhB*; coenzyme F420 hydrogenase subunit beta [EC:1.12.98.1] | SO_2016: 1<br>SO_2017: 8 | Methane metabolism, Redox reactions (*e.g.*, hydrogen oxidation) |
| *hdrA2*; heterodisulfide reductase subunit A2 [EC:1.8.7.3 1.8.98.4 1.8.98.5 1.8.98.6] / 4Fe-4S binding domain | NEUS: 1<br>SO_2017: 1 | Methanogenesis |
| *mcrD*; methyl-coenzyme M reductase subunit D | CA:1 | Methanogenesis, anaerobic methane metabolism |
| *mddA*; methanethiol S-methyltransferase [EC:2.1.1.334] | CA:1,<br>SO_2017:1 | Methionine metabolism to dimethyl sulfide |
| * *phoD*; alkaline phosphatase D [EC:3.1.3.1] | CA:3<br>NEUS:1 | Alkaline phosphatase |
| *^*rbcL*; ribulose-bisphosphate carboxylase large chain [EC:4.1.1.39] | NEUS:1<br>SO_2016:1<br>SO_2017:3 | D-ribulose 1,5-bisphosphate carboxylation, carbon dioxide fixation |
| *soxX*; L-cysteine S-thiosulfotransferase [EC:2.8.5.2] | CA:1 | Thiosulfate oxidation |

The highest confidence AMGs from Table 6 were alpha-amylase, *frhB*, and *rbcL*, where each was flanked by viral genes. All other AMGs from Table 6 were present on elements predicted as viral by VirSorter2, were not the final genes on the scaffolds, and were flanked by short hypothetical proteins commonly annotated on viral genomes. Phylogenetic trees were used to place the alpha-amylase, *frhB*, and *rbcL* AMGs with their closest matches from archaea and bacteria, as well as within the context of well characterized reference sequences. The alpha amylase was near-identical to a hypothetical protein from a Prolixibacteriaceae bacterium within a cluster of glycoside hydrolase family 13 alpha amylase proteins, strengthening the prediction of its role in polysaccharide metabolism (Figure S3). The four high-confidence FrhB represented two different sequence types, both clustering within a clade of bacterial coenzyme F420 reducing hydrogenase beta subunits, indicating a likely role in redox cycling (Figure S4). The RbcL RuBisCO subunit clustered with a group of “RbcL-like” proteins, and was distinct from a previously reported virally-encoded RbcL (Figure S5) [47].

### Virally encoded CRISPR arrays

A total of 106,371 landfill-derived viral elements that were ≥5 kbp were included in examinations of virally encoded CRISPR arrays and novel Cas protein identification. Viral elements were not clustered with CD-HIT in this analysis to preserve differences in CRISPR array structure. Only 0.43% (454) of these viral elements encoded CRISPR arrays, a set which had an average length of ∼47 kbp. Of these, 9/454 CRISPR array-encoding viruses were targeted in the virus-host networks, by predicted hosts from seven different phyla. Viral arrays encoded 2-102 spacers with an average spacer count of 8, with 15 viral arrays encoding at least 30 spacers. Notably, the five longest CRISPR arrays were encoded on relatively short scaffolds (5,027-7,697bp) that did not contain any detectable viral genes according to CheckV, or any detectable host (bacterial/archaeal) genes. These five scaffolds were conservatively considered false positive viral elements, most probably deriving from other Mobile Genetic Elements (MGEs), such as plasmids.

The longest putative viral elements that encoded CRISPR arrays containing 30+ spacers included a 364 kbp jumbophage (55 spacers, longest virally-encoded CRISPR array in these data) and a 508 kbp megaphage (32 spacers), both of which are predicted to be complete viral genomes according to CheckV. Multiple CRISPR arrays were less common, with 48 viral elements predicted to encode two or more CRISPR arrays (10.6% of CRISPR-array-encoding viral elements), with a maximum of six predicted CRISPR arrays for a single element. In total, 519 CRISPR arrays were associated with the 454 array-encoding viral elements.

The viral element with six CRISPR arrays is a predicted jumbophage that is 421,673 bp in length and >95% complete, according to CheckV. All six arrays were confirmed using CRISPRCasTyper [48] and five of these arrays were confirmed with CRISPR-CasFinder [49], the latter of which assigns evidence levels of 1-4 (4 being the strongest) to its predictions. CRISPR-CasFinder predicted a total of 9 CRISPRs in this element, but 3/9 of those predictions were low confidence (evidence level 1) and were not identified by our workflow/CRISPRCasTyper, one had an evidence level of three and was not detected by our workflow/CRISPRCasTyper, and five had an evidence level of four and were detected both by our workflow and CRISPRCasTyper. CRISPRCasTyper also predicted a variety of CRISPR-Cas types and subtypes from this six-CRISPR-encoding viral element, which included the annotation of putative effector nucleases (Figure 3A). Three of the megaphages (with genome sizes of 678, 641, and 508 kbp) were predicted to encode CRISPR systems (Figure 3B). Two of these megaphages exceed the ∼620 kbp genome size of the previously largest circularized CRISPR-encoding phage [33].

**Figure 3.**
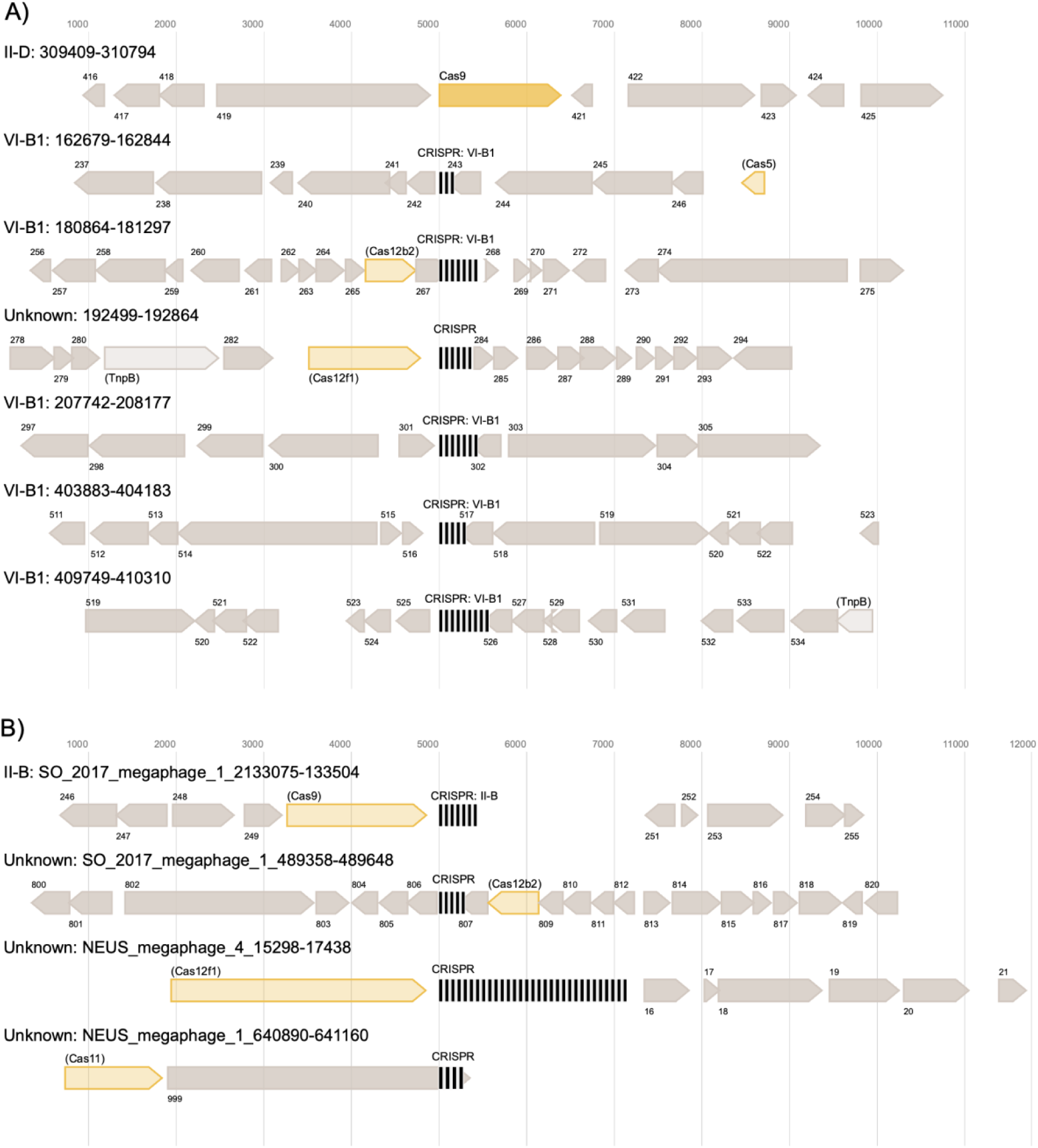
CRISPR in phage genomes of notable interest. A: CRISPR array and operons from a six-CRISPR-encoding viral element and B: CRISPR array and operons from all CRISPR-encoding megaphages. Cas protein interference/effector modules are noted in yellow and CRISPR arrays are denoted by clusters of vertical black lines. Unknown genes are dark grey, and putative TnpB homologs are light grey. All predicted interference/effector modules were considered speculative annotations [48] and are indicated by reduced opacity and parentheses around the predicted protein name. The most closely related Cas subtype classification is noted in each case, followed by the coordinates of each CRISPR array on each phage genome. ORFs proximal to predicted CRISPR arrays are numbered in order of occurrence within a single genetic element. In some cases, CRISPR arrays/CRISPR-Cas operons and their surrounding regions overlap with one another, resulting in ORFs depicted more than once in the figure. This image was generated using CRISPRCasTyper and processed using Affinity Designer.

### Virally encoded Class II effector nucleases

Viruses encode some of the most streamlined CRISPR-Cas systems identified to date [16,33,34]. Compact and efficient effector nucleases are targets of interest for biotech CRISPR-Cas applications. To identify putative streamlined Class II effector nucleases, a dataset of 1,494 ORFs ≥300 and ≤800 amino acids was identified from the 10 kbp regions upstream or downstream of the 519 putative virally encoded CRISPR arrays, including the five potentially non-viral MGEs. DNA-targeting class II effectors were predicted from these ORFs using profile HMMs and blastp searches.

For type V effector nucleases, a total of 161 putative type V nuclease ORFs were identified after dereplication. These ORFs were placed in a reference tree that included known type V nucleases and TnpB sequences (Figure 4). We observed no virally encoded nucleases that grouped with Cas12 subtypes a, b1, b2, c, d, e, f1, g, h, and i, likely because the majority of these protein families, with the exception of f1, b2, and g, have an average length that exceeds the 800 aa limit set for our ORFs of interest. The 800 aa limit was set to exclude Cas protein effectors that were not streamlined in size, i.e., not smaller in size relative to the currently most streamlined Cas effector, Cas12j/Cas12Φ, which is ∼800 aa in size ([16,34], see Methods for more details). We observed multiple nucleases that associated on the tree with other subtypes of Cas12 (i.e., f2 (1), f3 (1), j (1), and m (24)) and Cas14 (e (1), i (2), j (7), and k (9)), each of which average less than 800 aa in size per subtype. We examined each of these sequences for RuvC motifs, where some contained all expected RuvC motifs and others were less readily confirmed (see Supplemental Results for more information). We also observed two predicted TnpB sequences encoded by viruses. These putative TnpB sequences have all three RuvC motifs of TnpB and are 401aa and 433aa in length, respectively, in line with TnpB lengths of 350-550 aa [50]. These sequences are both encoded proximal to CRISPR arrays, consistent with the hypothesis that TnpB associated with CRISPR multiple times to give rise to different type V CRISPR-Cas systems, and the observation that TnpB is frequently encoded by phages [33]. In total, 114/454 (25%) of viral elements that encode CRISPR arrays were predicted to encode a type V homolog, consistent with the recently-observed enrichment of these hypercompact systems in phage [33] compared to their limited presence in bacterial and archaeal genomes [51]. Of these 114 viral elements, 10 (8.7%) are predicted to encode 2-3 unique type V effector proteins. These ten viral elements had an average length of 155,173 bp (15,118 – 421,673 bp) with 6 being larger than 120 kbp, whereas the average length of all viruses that encoded at least one type V system was 57,638 bp. One of these ten viral elements encodes three distinct predicted V-U4 nucleases, and all ten of these viral elements was predicted to encode at least one subtype V-U nuclease, or a nuclease that is most closely related to a V-U subtype nuclease. Lastly, we observed multiple nodes/clades on the Type V phylogeny that only include sequences from our study (Figure 4), representing additional unexplored novelty within this family that are obvious targets for nuclease activity characterization. None of the phages encoding the putative effectors highlighted on Figure 4 were predicted to encode the spacer acquisition machinery essential to adding new spacers to CRISPR arrays. This is consistent with previous studies showing that nearly all CRISPR-encoding phages lack adaptation machinery [16,33].

**Figure 4:**
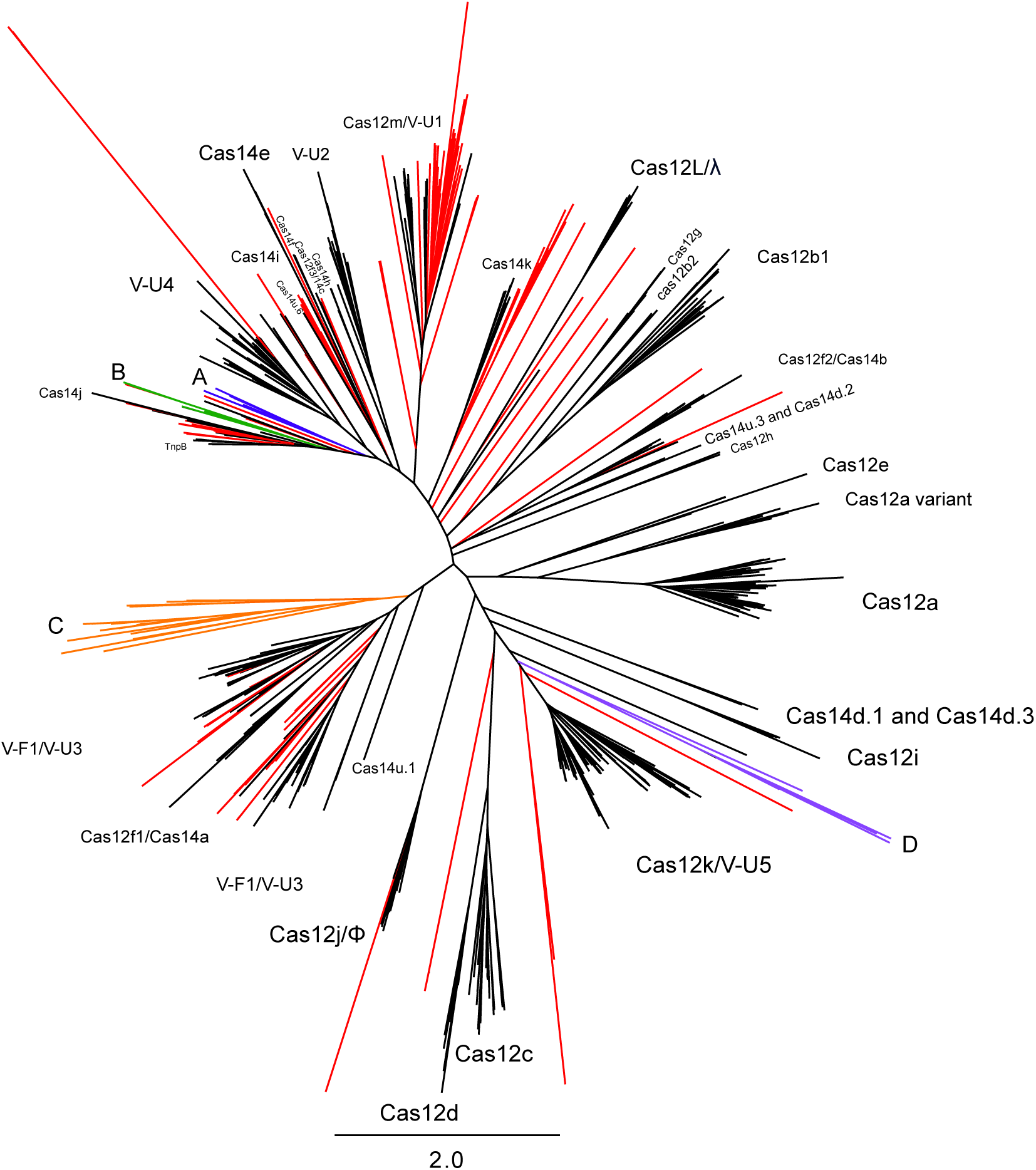
Type V CRISPR-associated nucleases, including virally encoded effectors from this study and reference nucleases. Putative nucleases from this study are coloured, where those in known groups or singletons are in red, and those in clades of further interest are denoted by a colour and a letter: A+blue, B+green, C+orange, and D+purple. Reference nucleases are in black. Names assigned to clades are those of (putative) effector nucleases, regardless of whether they have been experimentally characterized. In cases where the Type V subtype of a predicted nuclease has been classified, but the predicted nuclease has not been assigned an official Cas protein name, the predicted effector nucleases (e.g., c2c9; [51]), is denoted in the tree by its subtype classification (VU-2). Legacy names of nucleases that have been renamed during reclassifications are included for clarity, as are the subtypes of recently characterized nucleases, even if they have received an official Cas protein name as nuclease/subtype (e.g., Cas12m/V-U1). Singletons and landfill-exclusive clusters not associated with a specific named clade are unlabeled.

### Potentially novel Class II effector nucleases

All clades within the virally-encoded Type V effector phylogeny that included only our sequences (Figure 4) were examined on a case-by-case basis for motifs of interest. We identified 6 sequences (300-399 aa) that grouped proximal, but distinct to Cas14j (Figure 4, A/blue at 10 o’clock). These sequences had very weak RuvCI motifs, weak but stronger RuvCII motifs, and only the three longer sequences (385, 398, and 399 aa) had detectable RuvCIII motifs. A second cluster of 7 sequences (372-534 aa) was not associated with a specific Cas protein clade on the tree (Figure 4, B/green, 9 o’clock). All 6 had all 3 detectable RuvC motifs (i.e., RuvC[I,II,III]), but only two sequences had strong signatures of all RuvC motifs. We identified a clade of 19 sequences (Figure 4, C/orange, 8 o’clock) that, from sequence divergence, was predicted to be a new subtype of Type V effector protein, however, manual examination of these 19 sequences revealed no RuvC motifs. A final cluster of five sequences was identified proximal to the Cas12k/VU-5 clade (Figure 4, D/purple, 4 o’clock), but only one of these sequences had a single partial RuvC motif, specifically RuvCIII. Numerous singleton effectors were present on the tree, with most containing zero or only one identifiable RuvC motif (in all cases, RuvCIII).

## Discussion

Here we examined the diversity of viruses across 27 locations within three municipal waste sites. Viral DNA represented 2-13.5% of the landfill metagenomes, and depicted diverse viral communities. Limited overlap between viral communities from wastewater and groundwater environments with our landfill samples was observed (Figure S1). When we compared our landfill communities relative to each other using vConTACT2, we observed multiple Viral Clusters (VCs) that either included viral elements from more than one landfill site or were closely connected to VCs of viral elements from another landfill site (Figure S2). This suggests that the landfill virome is conserved, but distinct relative to the wastewater and groundwater viromes. Using geNomad, ∼97% of landfill viral elements were classified at the class level, but only ∼1.7% were classified at the order level (or family level, in cases where there was no assigned order) to a total of 15 orders and 12 families. The inability to classify most viral elements beyond Class is a clear indication of the lack of representation of these environmentally-derived viruses in current reference databases. This is further evidenced by the absence of viral RefSeq reference genomes in over 98% (6,023/6,116) of the VConTACT2 VCs that include our landfill viral elements.

Across all landfill datasets, we identified a total of 138 jumbophage genomes (i.e., those ≥200 kbp in size) with seven predicted to be complete or nearly complete megaphage genomes (i.e., ≥500 kbp). One of these putative megaphage genomes, SO_2017_megaphage_1, is ∼678 kbp long, which is, to our knowledge, the third largest phage genome reported to date. It is noteworthy that the two largest megaphage genomes reported in literature (735 and 717 kbp) were described from reports that examined 13 ecosystems [16], and 231 freshwater metagenomes [43], respectively – these reports were also published over three years apart. This suggests that bacteriophage genomes of this size are very rare in nature, making seven megaphage genomes from only two ecosystems (landfills and their adjacent groundwater wells) an unexpected discovery. The number of genes encoded by the seven megaphages ranges from 643 to 1,128, with 10-20 predicted AMGs per genome (Table S4). Three of the seven megaphages encode CRISPR arrays, with the 641 kbp and 678 kbp phage genomes now being the largest reported phage genomes to encode CRISPR arrays (exceeding the previous length of 620 kbp reported by [33]). None of the seven megaphages were targeted by MAGs within our virus-host networks, nor did the CRISPR arrays encoded by these viruses target any other viruses in our data. We were able to identify the large terminase subunit genes for six of the megaphages, which we used to place them in a reference tree of huge phages (Figure 1). Four of our megaphage genomes >600 kbp grouped together in a clade that included some the largest phage genomes described in literature [16,43], all of which are provisionally classified as “Mahaphages” and predicted to infect members of the phylum Bacteroidetes [16,43]. This suggests that while huge phages are taxonomically diverse [16], the phages with the largest genomes have shared ancestry across unconnected environments.

In our virus-host networks, as seen in our previous work [9], we observed hyper-targeted viral elements and viral elements predicted to infect across distinct phyla. Our consistent observation of putative cross-phylum infecting viruses in landfills, despite this phenomenon being reported from a limited number of other studies [1,8,10,11], suggests that landfills may be enriched for these broadly infecting viruses. Recent work showed that these very broad host range viruses are most often observed in syntrophic biofilms found in densely populated ecosystems [11]. Landfills are typically anaerobic systems where labile carbon is utilized fully through microbial handoffs and syntrophy, as evidenced by consistent and long-term methanogenesis [52]. Syntrophy is predicted to be widespread in landfills [52], and biofilm modes of life likely dominate in solid waste repositories, meaning the ecosystem characteristics of landfills are consistent with those predicted to select for broadly infecting viruses [11]. While we manually curated our putative cross-phylum interactions, and what we observe is consistent with existing literature, we note that because we solely utilize CRISPR spacer-viral protospacer matches to identify these interactions, these observations could stem from incorrectly binned CRISPR arrays. *In-vitro* evidence is necessary to substantiate these predictions.

We identified conserved CRISPR spacers across our different landfill samples to test whether landfills with higher similarity in their viral communities, and higher levels of virus-host cross-targeting between sites, also shared similar protospacer targets in viral genomes. Consistent with our examinations of viral community similarity and cross-targeting, both SO_2016 and SO_2017 shared >160 spacers with NEUS_2019, and SO_2016, SO_2017, and NEUS_2019 all shared <20 spacers with CA_2019 (Table 5). Omitting shared spacers between SO_2016 and SO_2017, as they are temporal datasets from the same landfill, ∼1-3% of spacers were shared between SO samples and NEUS_2019 with 100% identity and equal length. This suggests that landfills that share more similar viruses are more likely to share host CRISPR spacers, however, these shared spacers do not account for all cross-targeting observed between these sites.

From a total of 106,371 viral elements, 454 (0.43%) were predicted to encode CRISPR arrays, a subset that has an average length of ∼47 kbp compared to an average length of ∼13 kbp for viruses lacking CRISPR arrays. These observations are remarkably consistent with a previous study examining virally encoded CRISPR systems in which 0.4% of phages encoded CRISPR-Cas systems and had an average length of 52 kbp [33] . CRISPR-encoding viruses in this study did not show any infection bias to a particular host taxon.

A jumbophage with an approximately 421 kbp genome was predicted to encode six CRISPR arrays, five of which were annotated by two state-of-the-art CRISPR prediction tools [48,49]. According to CRISPRCasTyper, only four of these six CRISPR arrays were proximal (i.e., within 5 kbp upstream or downstream) to any *cas* gene or gene predicted to encode a TnpB-like protein. In total, this jumbophage contains predicted genes encoding two TnpB-like proteins, one Cas9, one Cas12b2, and one Cas12f1 (Figure 3A). The predicted *cas9* gene was not proximal to any CRISPR arrays (Figure 3A). The disjunction of arrays and *cas* genes is potentially due to the guide RNAs from the two orphan CRISPR arrays directing effectors encoded elsewhere in the viral genome, or in the host genome [53]. The orphan arrays could also be explained by the presence of proximal effector genes that are too divergent to be detected by current models of *cas* genes. The potential presence of multiple class II effector encoding genes in a single viral genome, along with TnpB encoding genes, which may themselves be hypercompact DNA targeting systems [54], could aid these elements defense against Cas protein inhibitory systems, such as anti-CRISPR [55]. Since CRISPR-Cas systems also have roles in processes such as gene expression and signal transduction [56], and viral CRISPR-Cas systems can target genetic systems of the host organism that regulate immunity [28] and host gene regulation [16], it is possible that the expansive CRISPR-Cas repertoire of viruses with large genomes may allow these phage to exert greater control over their host organism during infection, resulting in the creation of a “virocell” [57,58].

The average length of the ten viral elements encoding more than one predicted type V system was ∼155 kbp, which is nearly three times larger than the average length of all viral elements that encode at least one type V system (∼58 kbp) (Figure 5). Combined with the observation that the average length of CRISPR-encoding viruses was more than three times that of viruses lacking CRISPR, this suggests that the evolutionary costs of CRISPR-Cas systems [59] may be more manageable for viruses with larger genomes.

**Figure 5:**
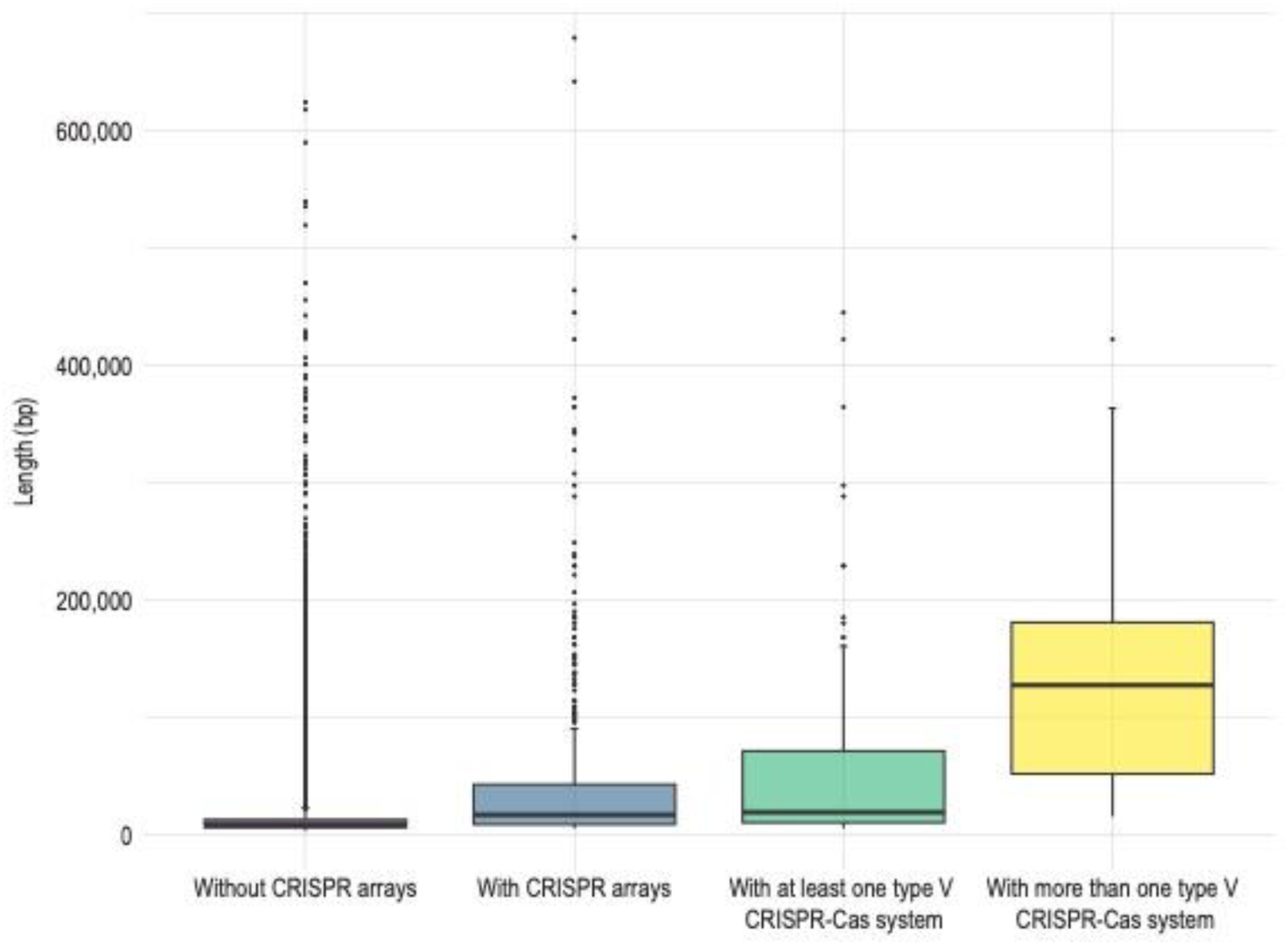
Boxplot of average viral element length associated with encoded CRISPR features. Boxes define the first to third quartile range with mean as the central bar. Outliers are reported as dots.

This manageable cost can be explained by jumbophages undergoing substantial coevolution of their capsid and genomes, where mutations that expand capsid volume in turn supports genome expansion [60]. Whether this capsid and genome coevolution extends beyond jumbophages and whether it can drive the acquisition of CRISPR-Cas systems remain open questions.

We detected virally encoded effector proteins that clustered with multiple subtypes of Cas12 (*i.e.,* f2, f3, j, and m), and Cas14 (e, i, j, and k), but most of our virally encoded type V-like effector proteins were located proximal to or with TnpB, the phage encoded systems Cas14j and Cas14k, and type V-U1, V-U5, and V-F1/VU-3 systems. All of these systems are either among the most streamlined type V or the progenitor (i.e., TnpB) systems. The subtype V-U systems are predicted as intermediates between TnpB and full-fledged type V effectors [51]. It is thus possible that viruses act as vectors for these intermediate systems, allowing an evolutionary path distinct from host-encoded systems, and with the potential to develop unique systems nearly exclusive to MGEs such as phages [16,33], and to MGE-associated systems such as Tn7-like transposons [31,61]. We identified 18 clades comprised of sequences exclusively from our study that were substantially divergent from previously described Type V effectors (Figure 4). Some of these proteins may represent novel type V nucleases, especially the seven for which we were able to identify all RuvC motifs. The full or partial absence of identified RuvC motifs in the other proteins could be due to substantial divergence of these motifs from those in reference enzymes, as is the case for phage-encoded Cas12L [33], or absence of canonical catalytic sites within the RuvC domain, as is the case for Cas12m [62]. Absent or divergent RuvC motifs do not necessarily indicate loss of function or alternate function for these Type V effector-related proteins. Even if these catalytic motifs are entirely absent, these proteins may still demonstrate interference activity through binding to, but not cleaving, their target nucleic acid, a function that has been seen for dCas9 [63] and Cas12m [62]. Both dCas9 and Cas12m have been repurposed as biotechnological tools, as alternate interactions with nucleic acids beyond targeted cleavage have downstream uses. For the putatively novel effector nucleases in this study to be considered as biotechnological tools, they would need to be validated *in vivo* for interference and/or cleavage activities.

## Conclusions

In summary, landfill viral diversity is relatively conserved, but distinct relevant to similar environment types such as wastewater and groundwater, and these viruses have rare or unique interactions with their hosts. Some of these viruses are predicted to broadly infect across the bacterial domain, which is consistent with recent, but limited literature. Bacteriophage genomes uncovered in this work are among the largest identified to date, sharing common evolutionary origins with the largest phage genomes described in literature. The discovery of seven of these phages from only 27 landfill metagenomes is in stark contrast to limited detection of these viruses even in studies that explore several hundred metagenomes [16,43]. Landfills may be a hot spot for discovering more of these megaphages, which may aid our understanding of the evolution of these phages and expand our knowledge of microbial genes that have been co-opted by giant phages. AMGs detected in this study have limited instances reported in literature, including genes involved in organochlorine and methane metabolism, both of which are relevant to landfills as contaminated sites that are globally significant producers of methane [36]. The discovery of AMGs in landfills that are relevant to landfill function but have rarely been observed elsewhere supports that genes mobilized within a microbial community are influenced by the selective pressures imposed by the environment.

We identified over 400 CRISPR-encoding viral elements from samples taken from three municipal landfills across North America. In some cases, these virally encoded CRISPRs were flanked by progenitor or full-fledged type V effector nucleases, including new branches of this protein family identified for the first time within this study. All of the putative type V effectors we identified in this study had a maximum length of 800 amino acids. If proven functional, these nucleases would help circumvent the size restrictions of certain viral delivery vectors such as those of Adeno-associated Viruses (AAVs) [64].

## Methods

### Sampling

Sampling of the Southern Ontario (SO) landfill occurred twice one year apart (July 2016 [SO 2016] and October 2017 [SO 2017]) and has been described previously [9,35]. Nine distinct cells of the NEUS landfill were sampled in February 2019 as described previously [36]. The final four metagenomes came from a closed landfill from Southern California (CA_2019) that was sampled in June of 2019. Leachate from one leachate well and the influent leachate to the leachate treatment plant were collected in sterile 500 mL containers. An additional sample of the treatment plant biofilter sludge was taken, with biomass split between planktonic and solid materials. All samples were passed through 0.2 µm Sterivex filters (Sigma Aldrich) until the filters clogged, with a minimum of three filters taken per sample (∼50-500 mL of volume processed per site). All landfill samples are detailed in Table S1.

The viral fraction of the SO landfill metagenomes was previously described from the perspective of viral-host interactions [9]. None of the other metagenomes have been analyzed for their viral fraction, and all cross-site comparisons as well as AMGs and megaphage analyses are novel to the current study. The CA_2019 landfill samples have not previously been reported.

### DNA extractions, sequencing, and host genome binning

DNA extractions for the SO landfill and adjacent aquifers and the NEUS samples have been previously described [9,35,36]. DNA extractions for the CA landfill were completed using the Qiagen Power Soil Pro kit following the manufacturer’s instructions with the addition of the sterivex filter directly into the bead beating tube in place of soil. The CA landfill samples underwent metagenomic sequencing at The Center for Applied Genomics (Toronto), generating 2x150 bp reads on an Illumina HiSeq. For metagenome assembly and host genome binning, the same pipeline as for previously analyzed landfill metagenomes was used. In brief, read trimming was accomplished with bbduk (https://github.com/BioInfoTools/BBMap/blob/master/sh/bbduk.sh) and sickle (https://github.com/najoshi/sickle). Scaffolds were assembled using Spades3 [65] (https://cab.spbu.ru/software/spades/) under the -meta flag with kmers 33, 55, 77, 99, and 127. Scaffolds ≥ 2.5 kbp were retained for further analyses. Read datasets were mapped to assembled metagenomes using Bowtie2 [66]. Scaffolds were binned using CONCOCT [67], MaxBin2 [68], MetaBAT2 [69], and consensus bins determined using DAS Tool [70]. Resultant bins were quality assessed using CheckM [71], and taxonomically classified using the Genome Taxonomy Database Toolkit [72]. Genome bins that were >70% complete and less than 10% contaminated (defined as, at minimum, medium quality draft Metagenome Assembled Genomes (MAGs; [73]) were used as putative host MAGs in further analyses. Metagenome sequencing, analysis, and host genome binning were previously described for the SO and NEUS samples [9,36] but follow this same workflow from read quality control onwards.

### Viral identification, classification, and genome binning

All scaffolds with a minimum length of 5 kbp were analyzed with VirSorter2 [37] and CheckV [38] sequentially, following the protocol at dx.doi.org/10.17504/protocols.io.bwm5pc86. Since CheckV, along with all other state-of-the-art tools, may not detect proviral region boundaries accurately, all proviruses identified by CheckV were discarded to maintain the integrity of further analyses, including AMG identification. All putative non-integrated viruses that had either at least one viral gene annotated by CheckV, or had zero viral and host genes annotated by CheckV, were retained for subsequent clustering with CD-HIT-est [39] using a global sequence identity threshold of c = 0.95. Viral genome binning was performed on the dereplicated set of viral scaffolds using vRhyme [40]. For viral taxonomic assignment, clustering, and comparisons of the landfill virome to related systems, predicted viral scaffolds with a minimum length of 10 kbp were input to vConTACT2 [44] along with 24,982 viral elements from wastewater samples and 20,541 viral elements from groundwater samples (all of which were ≥ 10 kbp and obtained from IMG/VR [45], and the vConTACT2 reference database “Prokaryotic Viral Refseq version 85 with ICTV-only taxonomy”). 10 kbp is the minimum size requirement for vConTACT2 hence viral scaffolds were size filtered accordingly. To assess the genetic similarity of landfill viruses across distinct landfills specifically, vConTACT2 was run a second time on all ≥ 10 kbp viral elements identified from our landfill metagenomes. Taxonomic classification of predicted viral elements >= 10kbp was performed using geNomad’s [74] annotate module and geNomad database v1.5.

In cases where we identified putative megaphages, we annotated terminase large subunit ORFs using HMMER3 [75] and Hidden Markov Models (HMMs) from the Virus Orthologous Groups database (https://vogdb.org/, release number vog218) and the Prokaryotic Virus Orthologous Groups (pVOGs) database [76]. Predicted large terminase subunits were confirmed with VIBRANT [46]. Predicted large terminase subunit sequences were aligned with other giant phage large terminase subunit sequences [16] using MUSCLE [77] version 5.1.linux64 and the Super5 algorithm. ModelTest-NG [78] was used to predict the best protein evolutionary model from the MUSCLE alignment. A phylogenetic tree was inferred using RAxML v.8.2.12 [79] under the PROTGAMMAVTF model and was visualized using Geneious [80].

### Virus-host linking

Viral elements that were at least 10 kbp in size were linked to their hosts through matching host CRISPR array spacers to viral protospacers. CRISPR arrays in host MAGs were detected using MINCED (https://github.com/ctSkennerton/minced) with the following parameters: -minSL [minimum spacer length] set to 25 and the -spacers flag included to output a fasta file of all identified CRISPR spacers. These CRISPR spacers were BLASTn [81] searched against viral elements with -task set as ‘blastn-short’, with alignments retained only if they had at most one mismatch, no gaps, a query coverage of ≥ 90% [2] and an E-value ≤ 10^-4^.

### Cross-phylum infection curation

All host MAGs involved in putative cross-phylum interactions were manually examined to assess the legitimacy of the interaction and to reduce the likelihood of incorrectly binned scaffolds leading to predicted cross-phylum interactions. First, the CRISPR-array encoding scaffolds of the MAGs predicted to target a putative cross-phylum infecting virus were ensured to be at least 10kb in length. Next, we required all MAGs predicted to be involved in cross-phylum interactions to have a maximum contamination of 5.00% according to CheckM.

### Assessment of CRISPR-Cas targeting space within and across landfills

Using the same workflow described above in “*Virus-host linking,*” we examined how often host CRISPR spacers from a site targeted viral elements from the same site and from different sites. We next detected spacers that were encoded multiple times in a single dataset or those that were shared between data sets, with no mismatches. Reverse complements were considered matches, but hits between a query spacer and a subsequence of a larger spacer were not considered as matches, as an exact length match requirement was implemented.

### Auxiliary metabolic gene (AMG) detection

AMGs were detected by inputting all dereplicated viral scaffolds into VIBRANT, which provides AMG predictions, with the ‘--virome’ flag. Predicted AMGs with limited or no prior representation in literature were considered high interest AMGs and manually curated as follows: the genomic neighborhoods of these putative AMGs were examined for obvious host genes, flanking viral hallmark genes, and hypothetical viral genes to assess whether predicted AMGs were due to microbial scaffolds misannotated as viral scaffolds or microbial genes proximal to integrated prophage. If a putative AMG of high interest had viral genes upstream and downstream of it or was encoded by a viral element that was predicted to be circular, it was considered a likely AMG. If a putative AMG of high interest only had viral genes on the upstream or downstream side and was not in a viral element predicted to be circular, it was considered a possible AMG. If a putative AMG of high interest did not have any viral genes directly upstream or downstream to it and was not encoded by a viral element predicted to be circular, it was discarded from subsequent examinations of high interest AMGs. To compare the incidence of high interest municipal waste site-derived AMGs to other systems, 100,000 viruses (≥ 5 kbp) were randomly sampled from IMG/VR v4 [82] and subjected to the same AMG detection pipeline.

Phylogenetic trees were made for virally-encoded alpha-amylase, coenzyme P450 hydrogenase subunit beta (FrhB), and RuBisCO large subunit (RbcL). For each protein, reference sequences were identified through best blastp hits (5-10 sequences selected, covering a broad taxonomic range), as well as from UniProtKB (FrhB, RbcL) or literature reviews of the protein family (alpha amylase). Alignments were generated using Muscle v. 3.8.425 [77] and masked to remove columns with more than 90% gaps in the Genious [80]. ModelTest-NG [78] was used to predict the best protein evolutionary model from the MUSCLE alignments. Phylogenetic trees were inferred using RAxML v.8.2.12 [79] and were visualized using Geneious.

### Viruses encoding CRISPR arrays

All predicted viral elements were screened for CRISPR arrays using MINCED (https://github.com/ctSkennerton/minced) with the following parameters: -minSL [minimum spacer length] set to 25 and the -spacers flag included to output a fasta file of all identified CRISPR spacers.

### Cas effector genes proximal to viral CRISPR arrays

CRISPRCasTyper [48] was used for automated detection of CRISPR-Cas operons in viral elements predicted to encode CRISPR arrays. Manual detection of Class II effector nucleases was performed as follows. All 10 kbp regions directly upstream or downstream from CRISPR arrays were examined for Open Reading Frames (ORFs) using prodigal [83] (flags -m -p meta -q). All predicted ORFs that were 300-800 amino acids (aa) in length were retained for further analyses to identify streamlined Class II proteins. This 800 aa limit was set to exclude Cas protein effectors that were not smaller in size than the current most streamlined Cas effector subtype, Cas12Φ/Cas12j, whose members are ∼800 aa in size [16,34]. Class II effector-encoding *cas* genes proximal to CRISPR arrays were predicted by querying CRISPR-array-adjacent ORFs using profile Hidden Markov Models (HMMs) [84] obtained from previous studies [48,85] and HMMER v.3.2.1 [75]. Additionally, BLASTp [81] searches were performed using multiple protein families that have been previously classified, and in some cases experimentally validated, as queries: TnpB (the predicted ancestor of Cas12 nucleases [51,86]); subtypes of Cas12; V-U nucleases (some of which have not been validated for interference activity); and Cas14 [16,51,62,87]; along with the recently identified and experimentally validated phage-encoded Cas12Φ (Cas12j) and Cas12λ (Cas12L). The screen did not include type VI effector proteins as queries, which are RNA-guided RNA targeting systems, as previous work demonstrates these systems aren’t nearly as common in viruses compared to the other Class II systems, i.e., type II and type V [33], and we were specifically interested in systems capable of DNA manipulation.

### Identification of putatively novel Cas protein effectors

Examinations were focused on type II and type V nucleases, as diverse nucleases of these types were previously identified in phages [33]. All ORFs that hit to type II nucleases were manually examined for the presence of RuvC and HNH motifs. ORF hits to type V nucleases were gathered. Due to some type V nucleases having very low identity to other subtypes [33], we also generated a separate dataset of all ORFs proximal to viral CRISPR arrays that were not identified as matches using our profile HMM or BLASTp searches (“Proximal ORFs”). The Proximal ORFs were combined with all reference type V nucleases and TnpB sequences, and aligned using MUSCLE version 5.1.linux64 and the super5 algorithm. ModelTest-NG [78] was used to predict the best protein evolutionary model from the MUSCLE alignment. A phylogenetic tree inferred using RAxML v.8.2.12 [79] with -m PROTGAMMAVTF was visualized using Geneious. Since the Proximal ORFs/proteins that were not hit by reference HMMs could include *cas* genes that don’t encode effector nucleases, or genes unrelated to CRISPR-Cas systems, putative effector proteins in this set were identified through BLASTp searching representatives of each clade of sequences on the phylogenetic tree against NCBI’s non redundant protein database (nr). A clade was retained only if the proteins/domains hit in nr were hypothetical proteins, TnpB, or endonucleases (excluding restriction endonucleases). Each retained clade was manually examined for RuvC motifs, and those with a detectable RuvC signature that also clustered close to known type V effector proteins and TnpB in the phylogenetic tree, were retained as putative Cas effector proteins. This subset of the Proximal ORFs dataset was then combined with the predicted effector ORFs that were identified through the HMM and/or BLASTp searches. This final aggregated dataset was aligned using MUSCLE version 5.1.linux64 and the PPP algorithm, due to smaller input size. ModelTest-NG [78] was used to predict the best protein evolutionary model from the MUSCLE alignment [77]. A phylogenetic tree was inferred using RAxML v.8.2.12 [79] with -m PROTGAMMAVTF and was visualized using Geneious (Figure 4).

## Declarations

### Ethics approval and consent to participate

Not applicable.

### Consent for publication

Not applicable.

### Availability of data and materials

The datasets generated and/or analysed during the current study are publicly available as follows:

*The SO_2016 metagenomes*: The raw reads are available in the SRA database under accessions SRS2844134 (CLC1_T1), SRS2845087 (CLC1_T2), SRS2844583 (LW1), SRS2844135 (LW2), SRS2844137 (LW3), and SRS2844136 (GW1). Assembled and annotated scaffolds are available on the Integrated Microbial Genomes and Microbiomes (IMG) database with the following IMG Genome IDs (Taxon Object IDs): 3300014203 (CLC1_T1), 3300014206 (CLC1_T2), 3300014204 (LW1), 3300015214 (LW2), 3300014205 (LW3), and 3300014208 (GW1) (https://img.jgi.doe.gov/cgi-bin/m/main.cgi). All MAGs from SO_2016 and SO_2017 connected in virus-host networks are available on NCBI under the BioProject PRJNA823399 and accessions JALNVE000000000-JALOCK000000000. Scaffolds predicted to be entirely or partially viral are available in IMG, with all viral IMG accessions listed in Supplemental File 1.

*The SO_2017 metagenomes*: The raw reads are available in the SRA database under accessions SRS4258091 (CLC), SRS12485771 (GW1), SRS4258089 (GW3), SRS12485774 (LW1), SRS4258090 (LW2), SRS12485775 (LW3), SRS4336822 (LW4), and SRS4258104 (SWC). The assembled and annotated SO_2017 metagenomes are deposited on IMG with the following IMG Genome IDs (Taxon Object IDs): 3300028602 (CLC), 3300037810 (GW1), 3300028028 (GW3), 3300035208 (LW1), 3300028603 (LW2), 3300037067 (LW3), 3300029288 (LW4), and 3300028032 (SWC). All MAGs from SO_2016 and SO_2017 connected in virus-host networks are available on NCBI under the BioProject PRJNA823399 and accessions JALNVE000000000-JALOCK000000000. Scaffolds predicted to be entirely or partially viral are available in IMG, with all viral IMG accessions listed in Supplemental File 1.

*The NEUS metagenomes* are housed under BioProject PRJNA900590. Within this BioProject, the raw reads files are available in the SRA database, under Biosamples SAMN31696084 – SAMN31696092. The 1,892 MAGs are available in the WGS database under accessions SAMN32731718 – SAMN32731810 (STA), SAMN32731811 – SAMN32731998 (STB), SAMN32733587 – SAMN32733720 (STC), SAMN32734194 – SAMN32734413 (STD1), SAMN32734415 – SAMN32734683 (STD2), SAMN32734737 – SAMN32734946 (STE), SAMN32737191 – SAMN32737484 (STF1), SAMN32737485 – SAMN32737723 (STF2). Viral scaffolds are available in the WGS database under accessions PP359733 - PP366656 and PP368833 - PP372551.

*The CA_2019 metagenomes* are housed under BioProject PRJNA1070006. The reads are available in the SRA database under accessions SRR27751815-SRR27751818. The MAGs implicated in viral-host interactions have been submitted to the WGS database (still processing, open science framework files available at DOI 10.17605/OSF.IO/FQ46R). Viral scaffolds are available in the WGS database under accessions PP366657 - PP368832.

### Authors’ contributions

NAG contributed to study conception and design, data collection, analysis and interpretation of results, draft manuscript preparation, and manuscript revision.

ZZ contributed to analysis and interpretation of results and manuscript revision.

KA contributed to analysis and interpretation of results and manuscript revision.

LAH contributed to study conception and design, data collection, analysis and interpretation of results, manuscript revision, and secured funding for the study.

## Acknowledgements

We are sincerely grateful to the landfill site management entities and their contracted consulting companies (anonymity by request) for site access and aid with sampling. We thank Dr. Jennifer Biddle and her lab group for hosting our team for sample processing in 2019. We thank members of the Hug Research Group, past and present, for help with landfill sampling.

