## Supplemental materials for "Discarded diversity: Novel megaphages, auxiliary metabolic genes, and virally encoded CRISPR-Cas systems in landfills"

**Table of Contents:**

**Supplemental Results** p. 1-2
**Supplemental Tables** p. 3-5
**Supplemental Figures** p. 5-9

### Supplemental Results

*Similarity to previously identified virally encoded CRISPR-Cas systems*

We observed several instances where our predicted effector sequences were clustered with or were most closely related to previously identified virally encoded CRISPR-Cas systems (i.e., Cas14j (7), Cas14k (9), Cas14i (2), Cas12j (1), and Cas12L (1), Figure 4), of which only Cas12j and Cas12L have been experimentally validated for function (Pausch et al. 2020; Al-Shayeb et al. 2022). Each of our relevant sequences was examined for RuvC motifs and assessed for similarity to the virally encoded nuclease it clustered most closely with.

Seven sequences ranging from 373-441aa clustered with previously identified Cas14j sequences (378-451aa; Figure 4, 10 o'clock). Our sequences showed high sequence similarity to

Cas14j sequences and contained all three RuvC motifs (RuvCI-III). Six additional sequences clustered proximal but distinct to the Cas14J cluster and are described in the next section. We detected two sequences of lengths 402 and 509 aa that clustered with Cas14i (Figure 4, 11 o'clock). Both of these sequences as well as reference Cas14i proteins had detectable RuvCI and RuvCIII motifs but very weak, if detectable, RuvCII motifs. We detected one sequence clustering with Cas12j (Figure 4, 7 o'clock). This sequence lacked the RuvCIII motif, as did Cas12j6, one of the 10 reference Cas12j nucleases (Al-Shayeb et al. 2020). Our putative Cas12j sequence was also missing key residues in the RuvCII domain, which Cas12j6 lacked entirely, and key residues in the RuvCI motif, all of which were present in Cas12j6. Only three orthologs of Cas12j(1-3) have been experimentally confirmed for function (Pausch et al. 2020). The comparisons made to Cas12j6 add confidence to the assignment of our query sequence as a Cas12j ortholog, despite its lack of key catalytic residues. Our putative Cas12j is the shortest within the clade, at 346 aa compared to 441 (Cas12j6 from a giant phage (Al-Shayeb et al. 2020)) and 708-813aa for the remaining 9 Cas12j proteins. While our sequence branches within the Cas12j clade, its activity is less confidently predicted based on the aberrant characteristics described above. Notably, our putative Cas12j sequence is encoded by a predicted plasmid, the second time a Cas12j-like protein was identified on a plasmid (Pinilla-Redondo et al. 2022).

### Supplemental Tables and Figures

#### Table S1: Landfill sites and sampling details

| Site | Sample ID | Sample type | Metagenome size (Gbp) | BioSample Accession | SRA Accession |
|---|---|---|---|---|---|
| SO_2016 | LW1 | Leachate well | 26.58 | SAMN07630781 | SRX3574636 |
| | LW2 | Leachate well | 30.00 | SAMN07630782 | SRX3574178 |
| | LW3 | Leachate well | 29.98 | SAMN07630780 | SRX3574180 |
| | CLC1_T1 | Composite leachate cistern | 29.89 | SAMN07630778 | SRX3574177 |
| | CLC1_T2 | Composite leachate cistern | 28.16 | SAMN07630777 | SRX3575198 |
| | GW1 | Groundwater well | 25.58 | SAMN07630779 | SRX3574179 |
| SO_2017 | LW1 | Leachate well | 15.57 | SAMN27259107 | SRX14723681 |
| | LW2 | Leachate well | 40.52 | SAMN10350574 | SRX5256784 |
| | LW3 | Leachate well | 47.58 | SAMN27259106 | SRX14723680 |
| | LW4 | Leachate well | 38.75 | SAMN10863920 | SRX5344198 |
| | CLC | Composite leachate cistern | 38.72 | SAMN10350766 | SRX5256785 |
| | SWC | Storm water catchment | 21.34 | SAMN10350495 | SRX5256798 |
| | GW1 | Groundwater well | 51.22 | SAMN27259105 | SRX14723679 |
| | GW3 | Groundwater well | 18.76 | SAMN10350765 | SRX5256783 |
| NEUS | A | Leachate well | 53.99 | SAMN31696084 | SRX18288880 |
| | B | Leachate well | 47.93 | SAMN31696085 | SRX18288881 |
| | C | Leachate well | 56.03 | SAMN31696086 | SRX18288882 |
| | D1 | Leachate well | 52.88 | SAMN31696087 | SRX18288883 |
| | D2 | Leachate well | 48.60 | SAMN31696088 | SRX18288884 |
| | E | Leachate well | 37.64 | SAMN31696089 | SRX18288885 |
| | F1 | Leachate well | 57.67 | SAMN31696090 | SRX18288886 |
| | F2 | Leachate well | 50.17 | SAMN31696091 | SRX18288887 |
| | CSWMC | Composite leachate cistern | 54.34 | SAMN31696092 | SRX18288888 |
| CA_2019 | LW1 | Leachate well | 31.10 | SAMN39634476 | SRX23416964 |
| | CLC | Composite leachate cistern | 64.38 | SAMN39634477 | SRX23416965 |
| | TP_BF | Treatment plant biofilter - planktonic | 61.73 | SAMN39634478 | SRX23416966 |
| | TP_BS | Treatment plant biofilter - solids | 58.01 | SAMN39634479 | SRX23416967 |

**Table S2: Putative cross-phylum host-virus interactions.**

| Sample set | Putative hosts | Host MAG phylum (GTDB-tk) | Host completion, contamination (%) | # host spacer to viral protospacer matches | Predicted viral element |
|---|---|---|---|---|---|
| CA_2019 | TPIn_75<br>TPBF_198 | Desulfobacterota<br>Proteobacteria | 99.41, 0.00<br>90.75, 0.63 | 1<br>7 | vMAG_518 |
| NEUS_ 2019 | STF2_137<br>STCSWMC_88 | Bacteroidota<br>Firmicutes_A | 95.56, 3.26<br>80.02, 4.08 | 1<br>2 | vMAG_1257 |
| NEUS_ 2019 | STCSWMC_93<br>STF1_64<br>STF2_19<br>STD2_245<br>STCSWMC_50<br>STCSWMC_25 | Bacteroidota<br>Bacteroidota<br>Bacteroidota<br>Bacteroidota<br>Cloacimonadota<br>Firmicutes_B | 94.35, 0.27<br>95.43, 0.27<br>95.43, 0.00<br>88.71, 2.42<br>98.90, 2.20<br>90.15, 4.60 | 15<br>16<br>1<br>5<br>1<br>3 | vMAG_3146 |
| NEUS_ 2019 | STF2_137<br>STCSWMC_88 | Bacteroidota<br>Firmicutes_A | 95.56, 3.26<br>80.02, 4.08 | 1<br>2 | vMAG_910 |
| NEUS_ 2019 | STF2_144<br>STCSWMC_50<br>STF2_148<br>STCSWMC_25 | Cloacimonadota<br>Cloacimonadota<br>Cloacimonadota<br>Firmicutes_B | 100.00, 1.10<br>98.90, 2.20<br>95.54, 1.10<br>90.15, 4.60 | 1<br>2<br>2<br>2 | NODE_3233_length_30680_cov_384 .260596_Sandtown_F2||full |
| SO_2017 | LW2_137<br>LW2_139 | Muirbacteria<br>Patescibacteria | 93.26, 4.56<br>70.53, 3.61 | 2<br>4 | vMAG_2310 |

**Tables S3 and S4 are included as a single .xlsx file "Supplementary File 1.xlsx"**

**Table S3**: Predicted AMGs encoded across all datasets.

**Table S4**: AMGs encoded by megaphage genomes.

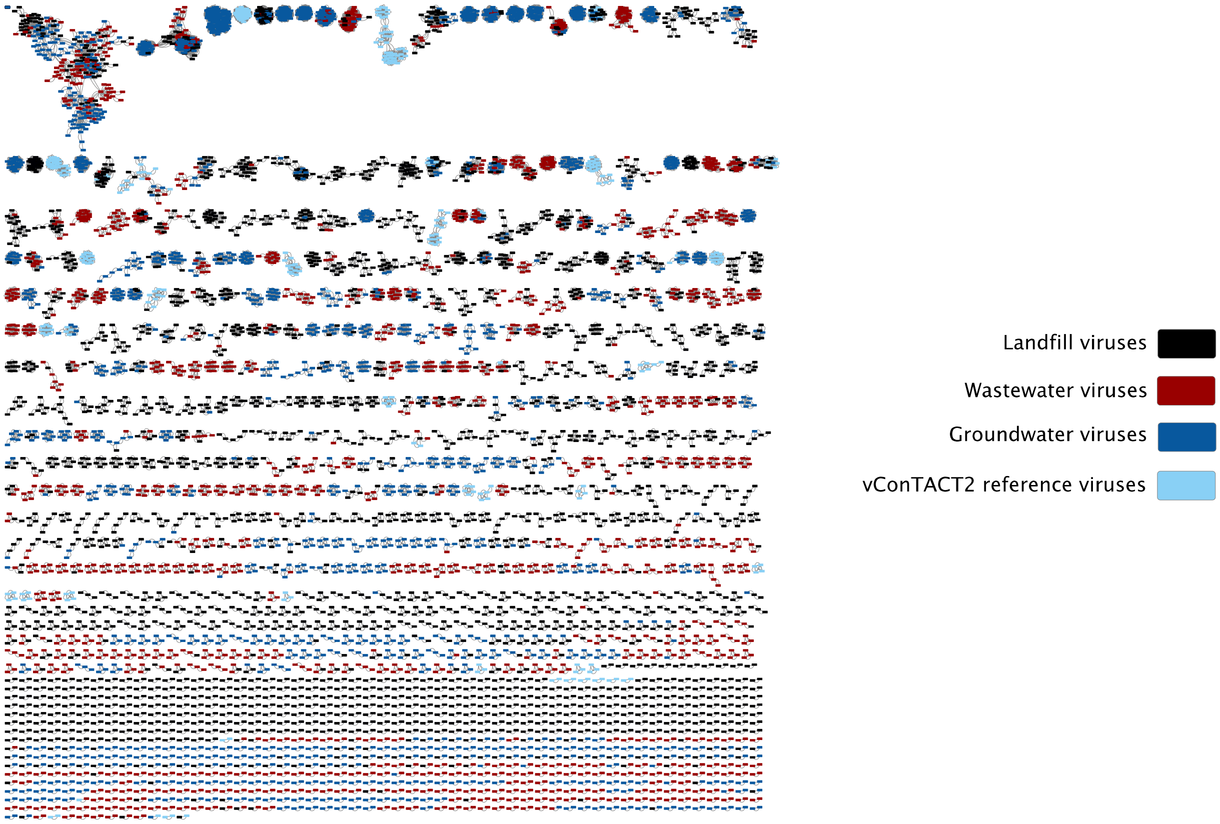


**Figure S1: Gene-sharing network of landfill viruses with related groups from IMG/VR.** Nodes represent viral elements and are coloured by sample site as summarized in the legend. Nodes connected by edges represent viral elements that share protein clusters. The network was generated using vConTACT2. The vConTACT2 reference database used was Prokaryotic Viral Refseq version 85 with ICTV-only taxonomy.

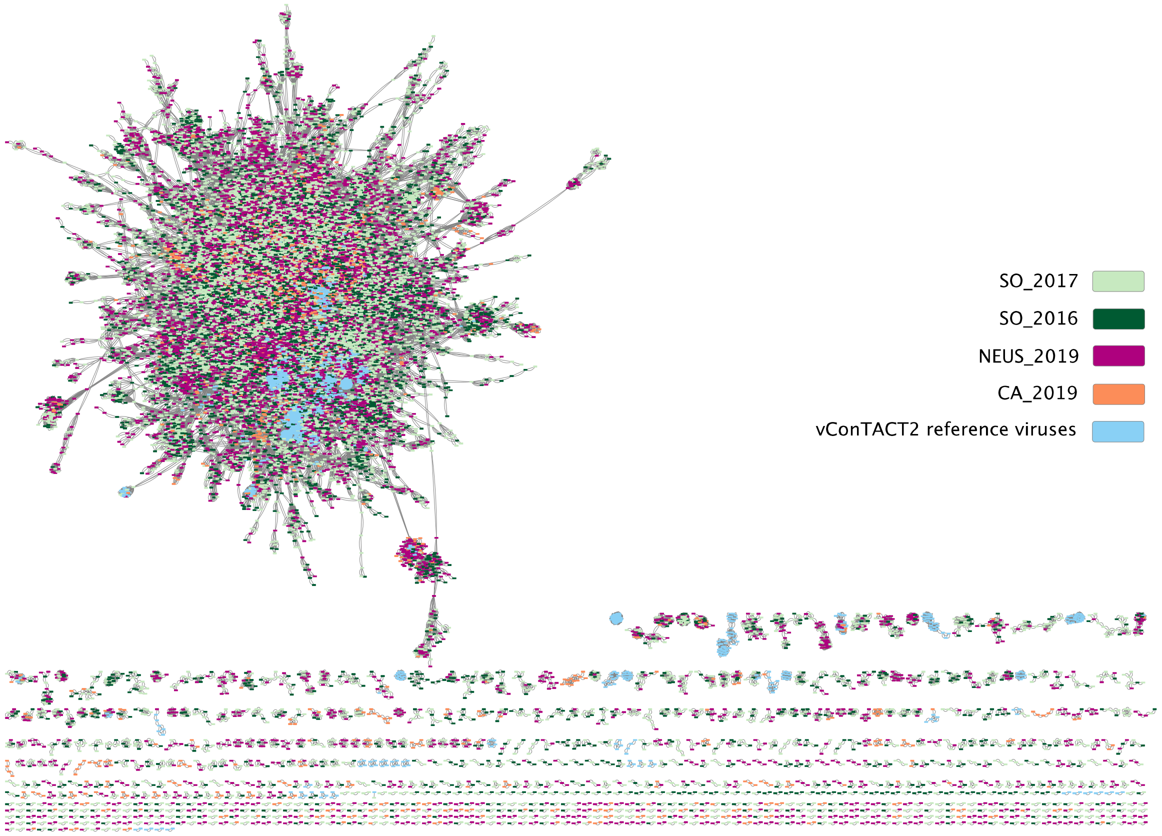


**Figure S2: Gene-sharing network of landfill viruses.** Viral nodes are coloured by the landfill they were identified in. Nodes connected by edges represent viral elements that share protein clusters. The network was generated using vConTACT2. The vConTACT2 reference database used was Prokaryotic Viral Refseq version 85 with ICTV-only taxonomy.

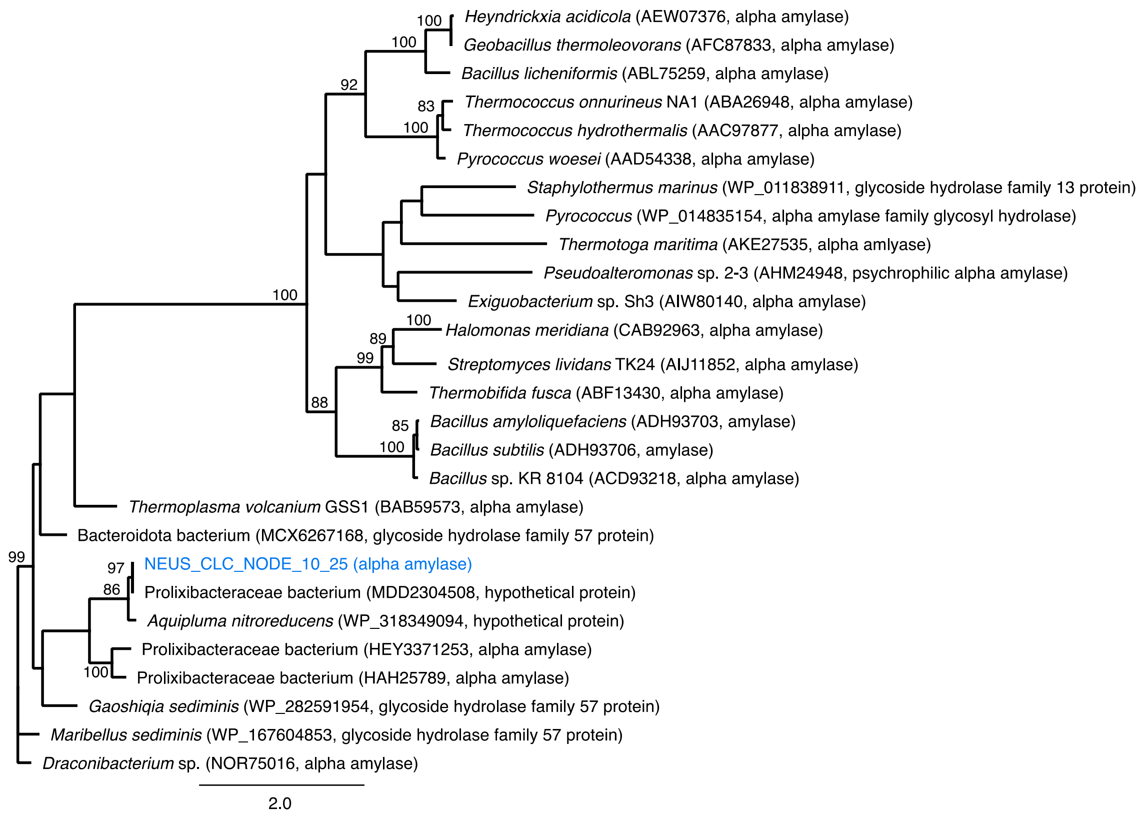


**Figure S3: A maximum likelihood phylogeny of the alpha amylase AMG (blue)**, including representative best hits from a blastp search and reference sequences (Mehta and Satyanarayana 2016). The final alignment contained 28 taxa and 805 unambiguously aligned columns. Alignments were generated with Muscle version 3.8.425 and (Edgar 2004) trimmed to remove columns with more than 90% gaps. The tree was generated using RAxML version 8 under the VT+I+G model of evolution (Stamatakis 2014) and visualized in Geneious (Kearse et al. 2012).

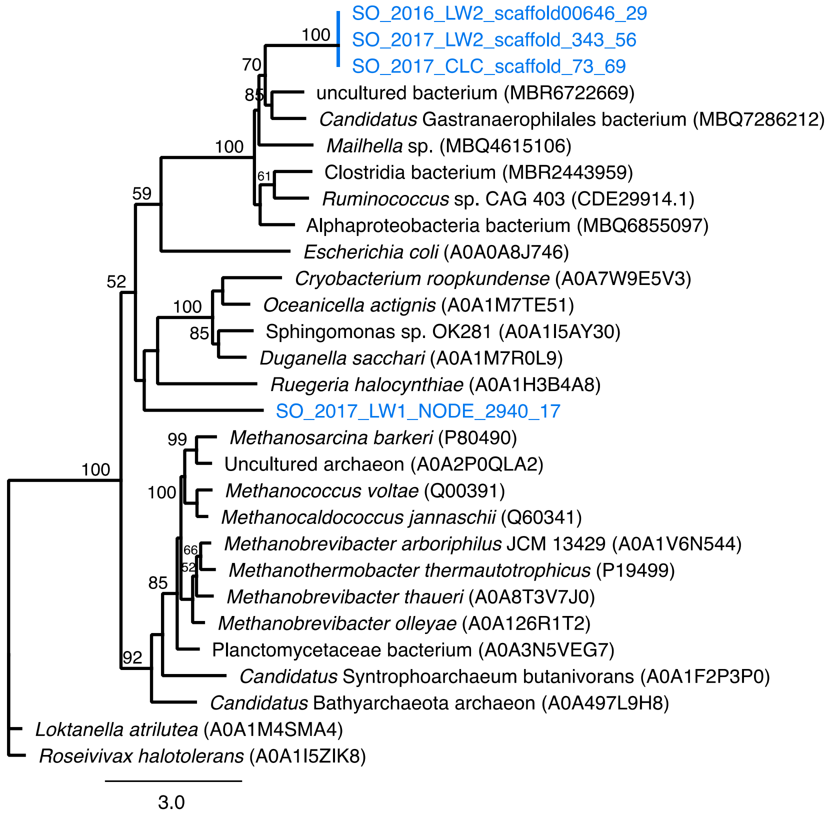


**Figure S4: A maximum likelihood phylogeny of the coenzyme P450 hydrolase subunit beta AMGs (blue)**, including representative best hits from a blastp search and reference sequences from UniProtKB. The final alignment contained 29 taxa and 845 unambiguously aligned columns. Alignments were generated with Muscle version 3.8.425 and (Edgar 2004) trimmed to remove columns with more than 90% gaps. The tree was generated using RAxML version 8 under the LG+I+G model of evolution (Stamatakis 2014) and visualized in Geneious (Kearse et al. 2012).

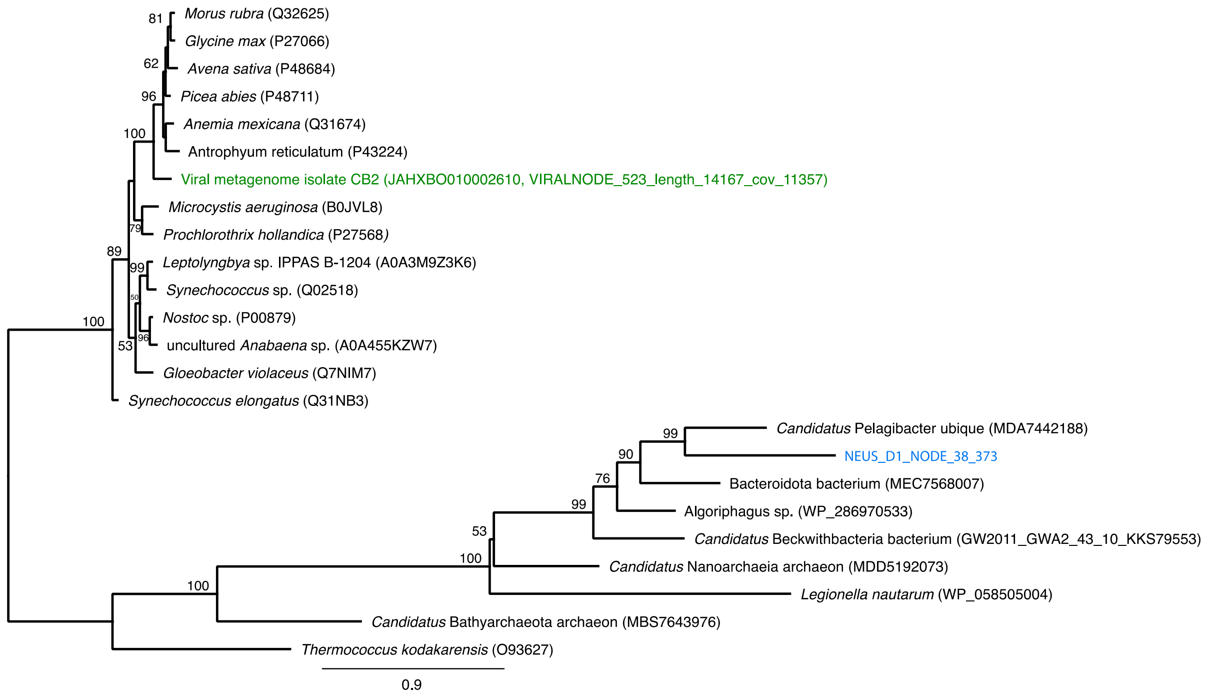


**Figure S5: A maximum likelihood phylogeny of the RuBisCO large subunit AMG (blue),** including representative best hits from a blastp search and reference sequences from UniProtKB. One previously reported RbcL from a viral fragment was also included (green, Bhattarai et al. 2021). The final alignment contained 24 taxa and 481 unambiguously aligned columns. Alignments were generated with Muscle version 3.8.425 and (Edgar 2004) trimmed to remove columns with more than 90% gaps. The tree was generated using RAxML version 8 under the LG+I+G model of evolution (Stamatakis 2014) and visualized in Geneious (Kearse et al. 2012).
